# Selective Sensitivity of Ph-like B-ALL to BRG1 Inhibition Reveals a Novel Targeted Therapy Strategy

**DOI:** 10.1101/2025.07.08.661805

**Authors:** V. S. S. Abhinav Ayyadevara, Shikha Gaur, Ashley Paik, Ria Perencsik, Monika Toma, Gerald Wertheim, Huimin Geng, Tomasz Skorski, Srividya Swaminathan, Christian Hurtz

## Abstract

Despite therapeutic advances, high-risk subtypes of B-cell acute lymphoblastic leukemia (B-ALL) such as Philadelphia chromosome-like (Ph-like) and *KMT2A*-rearranged (*KMT2A*-R) remain a formidable clinical challenge. BRG1 (gene name *SMARCA4*), the ATPase subunit of the SWI/SNF chromatin-remodeling complex, has been extensively studied in solid tumors, where inactivating mutations are linked to aggressive disease and poor prognosis. Although BRG1 is known to be essential for early B cell development, its role in B-ALL remains poorly understood. Therefore, we investigated the therapeutic relevance of BRG1 in high-risk B-ALL. Meta-analysis of gene expression data revealed that BRG1-inactivating mutations are exceedingly rare (0.35%) in B-ALL, suggesting that intact BRG1 function may be critical for leukemogenesis. Subtype-specific analyses revealed that elevated BRG1 expression is associated with significantly shorter overall survival in children with Ph-like B-ALL, while the opposite trend was observed in *KMT2A*-R B-ALL. We confirmed higher BRG1 expressions in Ph-like compared to *KMT2A*-R B-ALL via gene expression analysis, RT-PCR, and Western blotting. The pharmacologic inhibition of BRG1 using two selective inhibitors, BRM014 and FHD-286, revealed marked sensitivity in Ph-like B-ALL cell lines, whereas *KMT2A*-R B-ALL was resistant. Mechanistically, we found that BRG1 inhibition results in cell cycle arrest via downregulation of cell cycle regulators (*CCND3, CDK4, CDK6, E2F1*, and *MYC) and* upregulation of the cell cycle inhibitor *CDKN1B* (p27). Importantly, treatment with FHD-286 significantly prolonged the survival of *NSG* mice engrafted with Ph-like B-ALL cells. Taken together, these findings establish BRG1 as a critical, subtype-specific dependency in Ph-like B-ALL and demonstrate that its pharmacologic inhibition effectively suppresses leukemic cell proliferation through induction of cell cycle arrest. The pronounced *in vitro* sensitivity and improved *in vivo* survival upon BRG1 inhibition provide compelling preclinical evidence for its therapeutic targeting. These results support the advancement of BRG1-directed strategies as a viable treatment approach for patients with Ph-like B-ALL, with the potential to improve outcomes in this high-risk population.

**Highlights:**

1. Higher levels of BRG1 correlate to poor clinical outcomes in Ph-like but *KMT2A*-R B-ALL
2. Inhibition of BRG1 induces cell cycle arrest in Ph-like cells *in vitro* and extends the survival of mice in pre-clinical *in vivo* studies

*Graphical Abstract:* 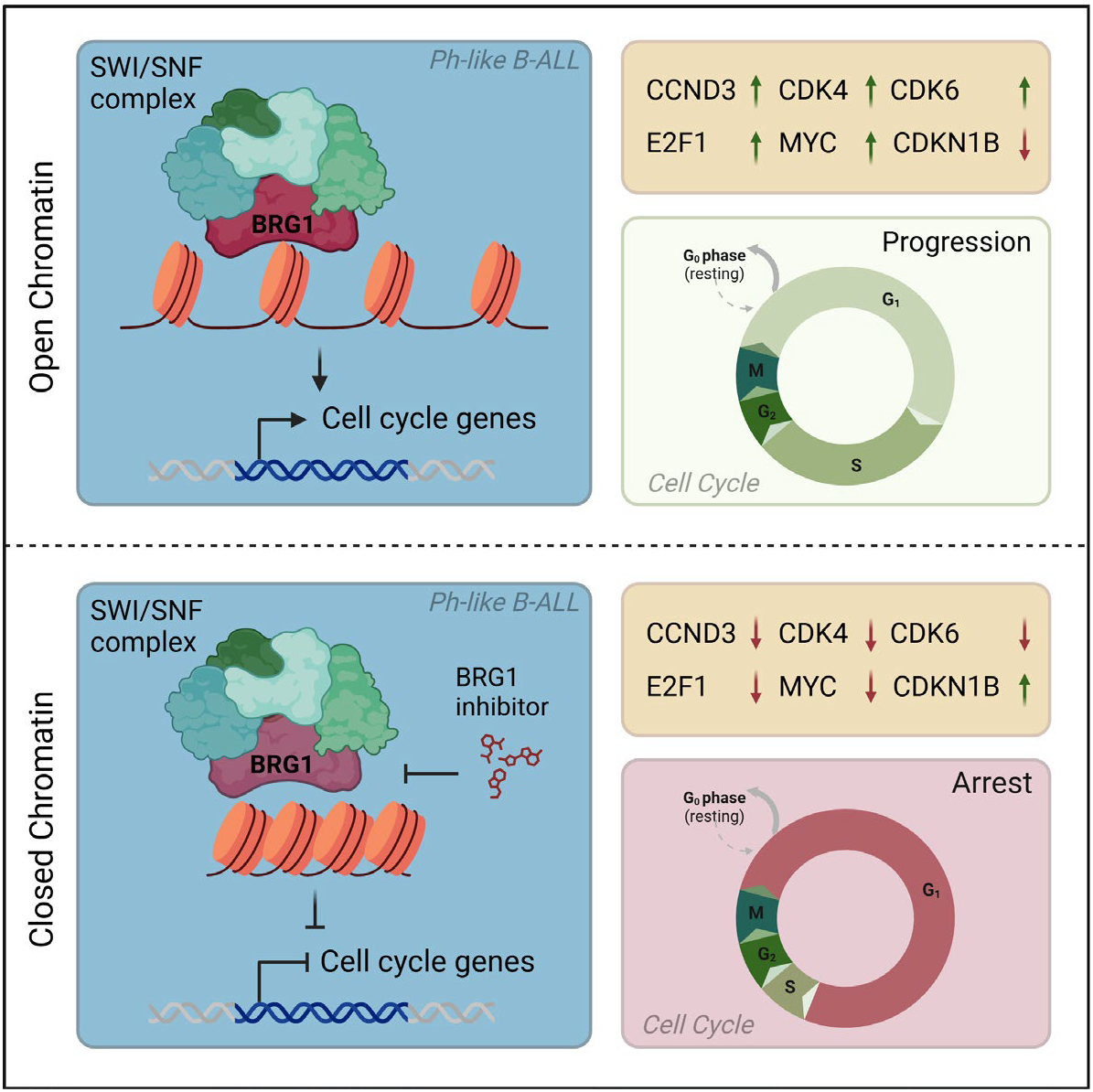

## Introduction

The Philadelphia chromosome–like (Ph-like) subtype of B-cell acute lymphoblastic leukemia (B-ALL) accounts for 10–30% of pediatric and adolescent/young adult (AYA) cases^1-3^ and up to 40% of adult B-ALL^3-10^. Despite therapeutic advances, patients with Ph-like ALL have poor outcomes, with high relapse rates and limited response to conventional chemotherapy^2,3,8,11^. This highlights the urgent need for alternative treatment strategies that target unique vulnerabilities specific to B-cell lineage and molecular subtype.

*SMARCA4*, encoding the BRG1 ATPase subunit of the SWI/SNF chromatin-remodeling complex, plays a context-dependent role in cancer^12,13^. In solid tumors such as non–small cell lung cancer, ovarian cancer, and rhabdoid tumors, *SMARCA4* is frequently mutated or deleted, supporting its role as a tumor suppressor. In contrast, in hematologic malignancies like acute myeloid leukemia (AML)^14^, BRG1 is essential, and its loss leads to a compensatory dependence on its paralog *SMARCA2* which encodes BRM^15^.

This synthetic lethality has spurred the development of *SMARCA2*-selective inhibitors. However, due to the conserved ATPase domain, most small-molecule inhibitors of the SMARCA family target both BRM and BRG1^15,16^. In stark contrast to solid tumors, *SMARCA4* is rarely mutated in B-ALL (<0.35%), suggesting that it is essential for leukemic cell function. Moreover, BRM is dispensable in B-ALL, further underscoring a lineage-specific dependency on BRG1.

Here, we investigated the functional requirement for BRG1 in B-ALL, with a particular focus on high-risk subtypes. Specifically, we (1) assessed the dependency of B-ALL on BRG1 using genetic and pharmacologic approaches, (2) identified subtype-specific expression and dependency patterns, (3) characterized the effects of BRG1 inhibition on cell proliferation and survival *in vitro*, and (4) evaluated the *in vivo* efficacy of BRG1 inhibition in preclinical models. These findings nominate BRG1 as a promising therapeutic target in Ph-like B-ALL.

## Results

### B-ALL Depends on Intact BRG1 and is Highly Expressed in Ph-like Subtype

B-cell development proceeds through a series of well-defined stages, beginning with common lymphoid progenitors (CLPs) and culminating in mature B cells. Each developmental step requires the activation of specific genes and transcriptional regulators to drive progression. The ATPase subunit *SMARCA4* (also known as BRG1) a core component of the SWI/SNF chromatin-remodeling complex, has been shown to play a critical role in early B cell development^17,18^. To evaluate BRG1’s role across B maturation, we performed a meta-analysis of *Smarca4* expression using publicly available gene expression dataset^19^. In one dataset profiling murine bone marrow subsets from CLP to Hardy fraction F, *Smarca4* mRNA expression was significantly elevated in fractions B, C, and D, corresponding to early pro-, late pro-/large pre-, and small pre-B cells^20^, respectively (**Figure 1a**).

**Figure 1:**
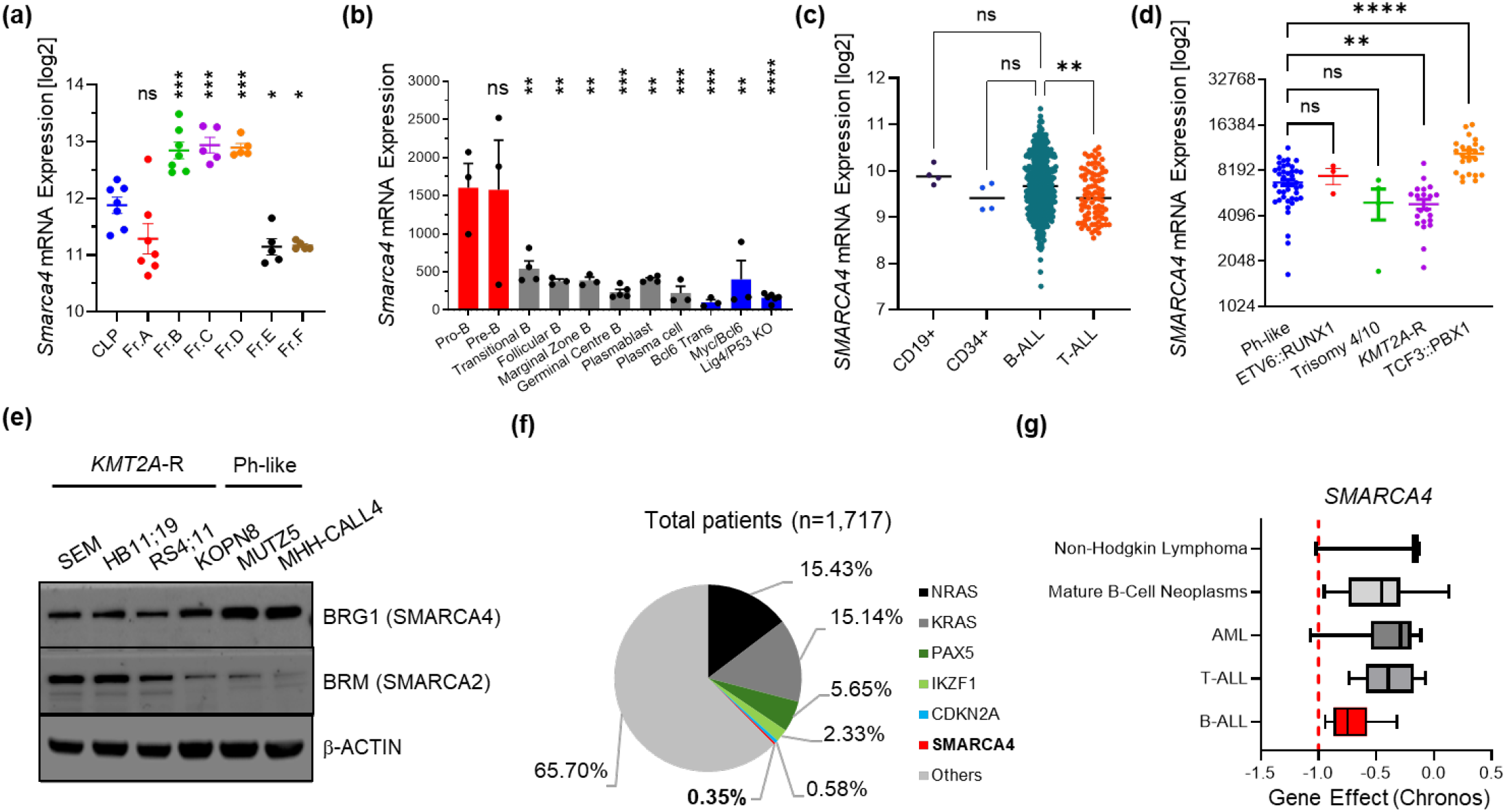
B-ALL Depends on Intact SMARCA4 With Some Subtypes Having Higher Expressions. **A)** *Smarca4* expressions in different murine hardy B fractions – ‘CLP’: Common lymphoid progenitor, ‘Fr. A’: Pre-Pro B, ‘Fr. B’: Early Pro B, ‘Fr. C’: Late Pro B / Large Pre-B, ‘Fr. D’: Small Pre-B, ‘Fr. E’: Immature B, and ‘Fr. F’: Mature B cells (GSE38463). **B)** Expression of *Smarca4* in different murine B developmental stages and lymphoma models (GSE26408). **C)** *SMARCA4* expressions in B-ALL populations compared to T-ALL, CD19+, and CD34+ populations (GSE33315). **D)** *SMARCA4* expressions in different subtypes of B-ALL populations (GSE11877). **E)** Western blot analysis of BRG1, BRM, and β-actin in four *KMT2A*-R (SEM, HB11;19, RS4;11, KOPN8) and two Ph-like (MUTZ5, MHH-CALL4) B-ALL cell lines. **F)** Mutational frequencies of different oncogenes in B-ALL patient populations from the COG study P9906 (GSE11877). **G)** Dependency analysis of *SMARCA4* via the Cancer Dependency MAP website. Data are represented as individual values with mean ± SEM bars. *P < 0.05; **P < 0.01; ***P < 0.001; ****P < 0.0001 by One-way Anova Dunnett’s multiple comparison test.

We validated this finding via a second analysis^21^ that included both normal murine B-cell subsets and murine lymphoma models (**Figure 1b**). These results suggest that *Smarca4* is specifically required in early B-cell development (red bars) but not in mature B cells (grey bars) or mature B-cell-derived lymphomas (blue bars). Given that B cell-derived acute lymphoblastic leukemia (B-ALL) cells originate from early progenitor B cells, these findings raise the possibility that B-ALL cells may retain elevated *SMARCA4* expression levels and depend on it for function and survival.

To explore the relevance of *SMARCA4* in B-ALL, we first analyzed a gene expression dataset^22^ that included normal human and leukemic cells isolated from pediatric patients. This dataset comprised healthy CD19+ and CD34+ cells, as well as B- and T-lineage ALL samples. We found that *SMARCA4* expression levels vary widely across B-ALL cases but were generally higher compared to both normal CD34+ and T-ALL samples (**Figure 1c**).

To determine whether specific B-ALL subtypes exhibit differential *SMARCA4* expression, we next analyzed gene expression data from the COG P9906 study^23^ which includes molecular-defined subsets of pediatric B-ALL. Our analysis revealed that *SMARCA4* expression was highest in TCF3::PBX1 and Ph-like subtypes compared to other subtypes such as ETV6::RUNX1, Trisomy 4/10, and *KMT2A*-R B-ALL (**Figure 1d**).

Given the relatively favorable prognosis associated with ETV6::RUNX1, Trisomy 4/10, and TCF3::PBX1, compared to the Ph-like and *KMT2A*-R B-ALL subtypes, we focused our subsequent analyses on the role of SMARCA4 specifically in these two high-risk groups.

Via western blotting and RT-PCR, we confirmed that BRG1 protein and *SMARCA4* mRNA levels are higher in Ph-like B-ALL cell lines (MUTZ5, MHH-CALL4) compared to *KMT2A*-R B-ALL cell lines (SEM, HB11;19, RS4;11, KOPN8) (**Figure 1e and Supplementary Figure 1a**). In contrast, BRM (encoded by *SMARCA2*) protein levels were notably lower in Ph-like but higher in *KMT2A*-R B-ALL cell lines (**Figure 1e**). These findings were further assessed at the transcription level using RT-PCR, which showed similar *SMARCA2* mRNA levels across both subtypes, without a consistent difference. (**Supplementary Figure 1b**). Analysis of the COG P9906 gene expression dataset^23^ supported these observations and among the different B-ALL subtypes, there were no significant differences in *SMARCA2* expression (**Supplementary Figure 1c**). Taking together, these data suggest that the elevated BRM protein levels observed in *KMT2A*-R B-ALL may be regulated via post-translational modifications (PTMs), and not solely at the mRNA level. Given the expression patterns of BRG1 and BRM in these subtypes, we hypothesize that Ph-like B-ALL primarily relies on BRG1, while *KMT2A*-R B-ALL depends on both BRM and BRG1 for chromatin remodeling.

Mutations in *SMARCA4* are common in solid tumors such as non-small cell lung cancer (NSCLC), where BRG1 acts as a tumor suppressor and its loss promotes more aggressive disease progression^24-26^. To assess the mutation frequency of *SMARCA4* in leukemia, and to determine whether increased mutation rates in specific subsets of ALL might explain the lower BRG1 levels observed in *KMT2A*-R ALL (**Figure 1e**), we analyzed whole-genome sequencing (WGS) and whole-exome sequencing (WES) data from 1,717 B-ALL patients included in the St. Jude study^27^. However, our analysis revealed that non-synonymous *SMARCA4* mutations are exceedingly rare in B-ALL overall, occurring in only 0.35% of cases (n=6; **Figure 1f, Supplementary Table 1**), suggesting that intact BRG1 may be essential for leukemic pathogenesis and, in contrast to its role in solid tumors, does not function as a tumor suppressor in B-ALL.

To validate this dependency, we examined CRISPR-mediated *SMARCA4* deletion data from the Cancer Dependency Map (DepMAP) portal. Strikingly, B-ALL emerged as the only hematologic malignancy with a selective dependency on *SMARCA4* (**Figure 1g and Supplementary Figure 1d**), in contrast other hematologic malignancies, including T-ALL, AML, and lymphomas, showed minimal dependency and correspondingly lower *Smarca4* expression levels (**Figure 1b and 1g**). Additionally, B-ALL dependency on *SMARCA2* was similar to other hematologic cancers and NSCLC (**Supplementary Figure 1e, f**), further underscoring that the selective reliance of B-ALL on BRG1 over BRM.

### Higher BRG1 levels Correlate with Poor Clinical Outcomes in Ph-like B-ALL

To assess whether pharmacological inhibition of BRG1 may have therapeutic relevance in B-ALL, we analyzed patient outcome data from the COG P9906 study by stratifying patients based on *SMARCA4* expression levels, categorizing as ‘high’ or ‘low’ relative to the cohort’s median expression. Across the entire B-ALL cohort, no significant differences in overall survival were observed between the high and low *SMARCA4* expression groups (**Figure 2a**). However, distinct subtype-specific patterns emerged upon further stratification. In Ph-like ALL, higher *SMARCA4* expression was significantly associated with poor clinical outcomes, whereas lower expression correlated with improved survival (**Figure 2b**). Conversely, in *KMT2A*-R B-ALL, the opposite trend was observed: higher *SMARCA4* expression corresponded with better survival, while lower expression correlated with poor outcomes (**Figure 2c**). In contrast, higher *SMARCA2* expressions were generally associated with poor clinical outcomes across all B-ALL cases, including Ph-like and *KMT2A*-R B-ALL (**Supplementary Figure 2a-c**). These findings suggest that Ph-like B-ALL patients could benefit from therapeutic strategies targeting BRG1, while patients with *KMT2A*-R ALL may respond differently.

**Figure 2:**
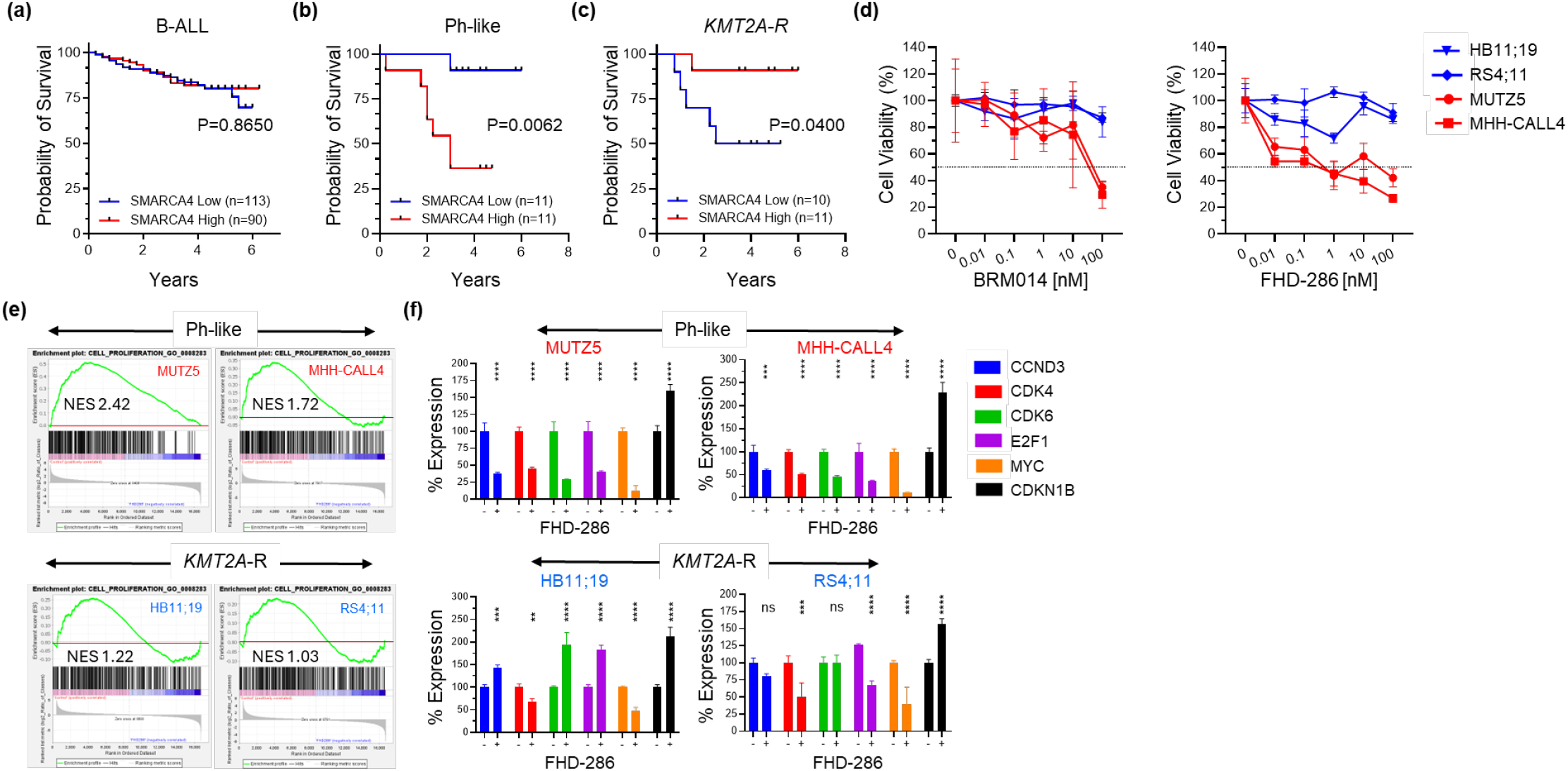
Ph-like B-ALL Cell Lines Are Sensitive to BRG1 Inhibition *In Vitro*. Kaplan-Meier analysis of the overall survival of all B-ALL **(A)**, Ph-like B-ALL **(B)**, and *KMT2A*-R B-ALL **(C)** patients from the COG study P9906 with high vs low *SMARCA4* mRNA expression (GSE11877). **D)** Ph-like (red; MUTZ5 and MHH-CALL4) and *KMT2A-R* B-ALL cell lines (blue; HB11;19 and KOPN8) were treated with increasing concentrations of BRM014 (left) or FHD-286 (right). Viability/cell proliferation was determined via an XTT assay after 72 hours. **E)** GSEA enrichment plots of Ph-like (MUTZ5 and MHH-CALL4) and *KMT2A-R* B-ALL cell lines (HB11;19 and KOPN8) after 24 hours of treatment either with control or 100 nM FHD-286. “NES” refers to the Normalized enrichment score (GSE297161). **F)** RT-PCR was performed to determine the gene expression levels of the indicated genes in Ph-like (MUTZ5 and MHH-CALL4) and *KMT2A-*R B-ALL cell lines (HB11;19 and KOPN8) after 24 hours of treatment either with control or 100 nM FHD-286. *ACTB* (Beta Actin) was used as a reference gene to calculate expression levels. Significance was calculated via a log-rank test in Prism **(A-C)**. Data are represented as individual values with mean ± SEM bars. *P < 0.05; **P < 0.01; ***P < 0.001; ****P < 0.0001 by Two-way Anova Sidak’s multiple comparison test **(F)** or two-sided Log-rank (Mantel-Cox) test **(A-C)**.

To functionally test this hypothesis, we utilized two recently developed dual BRG1/BRM ATPase inhibitors - BRM014^15^ and FHD-286^28^ - and assessed the sensitivity of Ph-like and *KMT2A*-R B-ALL cell lines to pharmacologic inhibition.

### Ph-like Cells are Sensitive to Pharmacologic Inhibition of BRG1/BRM in vitro

To evaluate the therapeutic potential of BRG1/BRM inhibition in B-ALL, we initially performed XTT cytotoxicity assays on two Ph-like and two *KMT2A*-R B-ALL cell lines *in vitro*. As predicted, Ph-like cell lines demonstrated high sensitivity to both BRM014 and FHD-286, with IC^50^ values of ∼50 nM and 1 nM, respectively, while *KMT2A*-R cell lines exhibited relative resistance to both compounds (**Figure 2d**).

To understand the mechanism underlying BRG1/BRM inhibition, we performed an RNA-seq experiment to identify differentially expressed genes (DEGs) in control vs FHD-286 treated Ph-like and *KMT2A*-R B-ALL cell lines. Gene Set Enrichment Analysis (GSEA) revealed that genes involved in cell proliferation are strongly enriched in control samples relative to FHD-286 treated cells (**Figure 2e**). This enrichment, measured by NES (Normalized enrichment score), was more pronounced in Ph-like B-ALL cell lines (MUTZ5, MHH-CALL4) compared to *KMT2A*-R B-ALL cell lines (HB11;19, RS4;11; **Figure 2e**).

We further validated these findings via RT-PCR, confirming that key regulators of cell cycle progression, including *CCND3, CDK4, CDK6, E2F1*, and *MYC*, were downregulated following BRG1 inhibition, while the cell cycle inhibitor *CDKN1B* (p27) was upregulated (**Figure 2f)**. Western blot analysis corroborated MYC protein downregulation in both Ph-like and *KMT2A*-R B-ALL cell lines treated with BRM014 or FHD-286, consistent with previous reports that used genetic approaches to inhibit *SMARCA4* expression^14,17^ (**Supplementary Figure 2d**).

To determine whether this dysregulation of cell cycle genes causes cell cycle arrest, we performed flow cytometry-based cell cycle analysis (BrDU). Treatment with BRM014 or FHD-286 resulted in a marked reduction of S-phase cells and accumulation in the G0/G1 phase across all cell lines. Notably, Ph-like B-ALL cells (MUTZ5, MHH-CALL4) exhibited a nearly complete loss of S-phase cells, indicating a profound cell cycle arrest. In contrast, *KMT2A*-R B-ALL cells (HB11;19, RS4;11) retained a substantial fraction of cells in S-phase, suggesting continued proliferation despite treatment (**Figure 3a**). Among the inhibitors, FHD-286 demonstrated relatively higher potency in inducing this arrest (**Figure 3a**).

**Figure 3:**
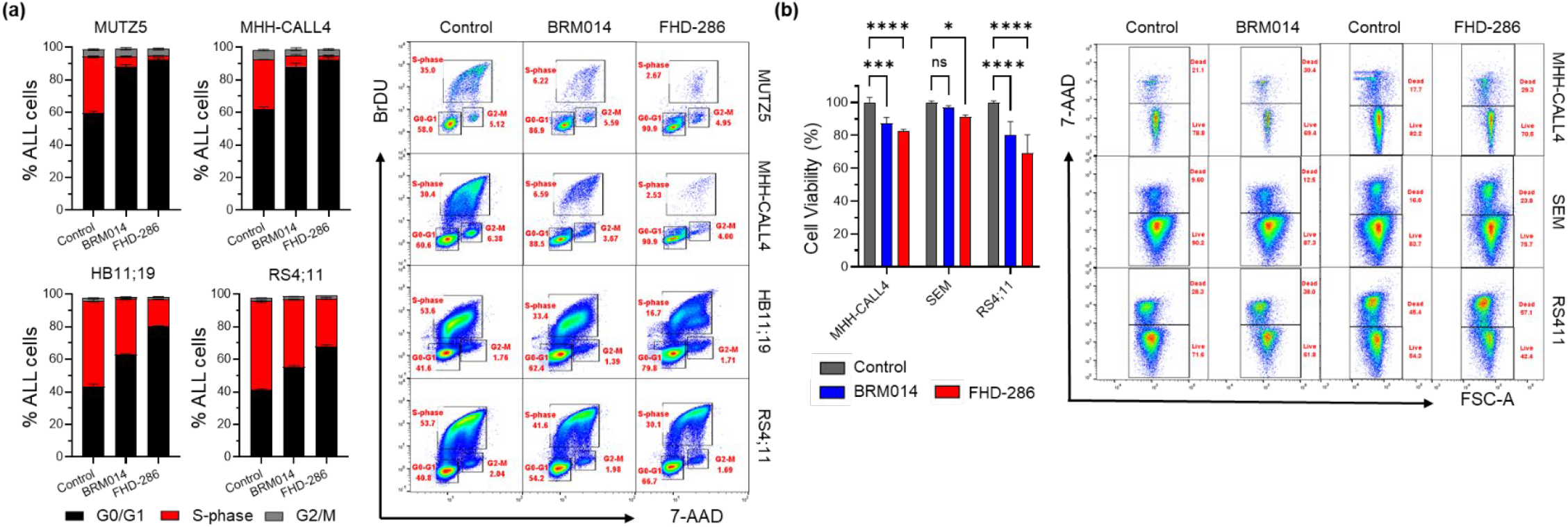
SMARCA4 Inhibition Results in Cell Cycle Arrest in Ph-like B-ALL *In Vitro*. **A)** Cell cycle analysis of the Ph-like (MUTZ5 and MHH-CALL4) and *KMT2A-*R B-ALL cell lines (HB11;19 and KOPN8) after treatment with 100 nM BRM014 or FHD-286 for 72h. On the left are representative examples of the flow cytometric analysis (n=3), and on the right is the summary of all three experiments. **B)** Cell viability and apoptosis were tested in control- and BRG1/BRM inhibitor-treated (BRM014 or FHD-286; 100 nM; 72h) Ph-like (MHH-CALL4) and *KMT2A*-R (SEM and RS4;11) B-ALL cell lines. On the left are representative examples of the flow cytometric analysis (n=3) and on the right is the summary of all three experiments. Data are represented as individual values with mean ± SEM bars. *P < 0.05; **P < 0.01; ***P < 0.001; ****P < 0.0001 by Two-way Anova Dunnett’s multiple comparison test.

To assess whether BRG1/BRM inhibition also triggers apoptosis, we conducted flow cytometry-based apoptosis assays (Annexin-V/7-AAD). These assays revealed that neither Ph-like nor *KMT2A*-R B-ALL cells undergo apoptosis following treatment (**Figure 3b**). To further investigate this, we used a human apoptosis protein array on Ph-like cell line due to its higher sensitivity to FHD-286 and found upregulation of anti-apoptotic proteins such as Survivin (*BIRC5*) and XIAP (X-linked inhibitor of apoptosis), along with downregulation of HIF-1α (Hypoxia-Inducible Factor 1-Alpha). Notably, we observed no activation of caspase-3 or p53 within 24 hours of FHD-286 treatment in the MUTZ5 cell line (**Supplementary Figure 2e**), suggesting a potential resistance mechanism to apoptosis.

Altogether, our *in vitro* results demonstrate that BRG1/BRM inhibition in Ph-like B-ALL primarily induces cell cycle arrest, rather than apoptosis, via transcriptional downregulation of cell cycle promoting genes and upregulation of cell cycle inhibitors.

### Pharmacologic Inhibition of BRG1/BRM Extends the Survival of Mice Engrafted with Ph-like B-ALL

To validate the *in vivo* therapeutic potential of BRG1/BRM inhibition to treat Ph-like B-ALL, we performed a xenograft study using *NSG* mice engrafted with the Ph-like B-ALL cell line MUTZ5. Mice were treated with either vehicle or the dual BRG1/BRM inhibitor FHD-286 (see Methods, **Figure 4a**). Kaplan-Meier survival analysis demonstrates a significant extension in median survival in the FHD-286-treated group (**Figure 4b**). This survival benefit was accompanied by slower body weight loss in the treatment group, indicating delayed disease progression and drug tolerability (**Figure 4c**). Furthermore, the FHD-286 group had to be sacrificed early because of weight loss, not because of progressive disease. Had this weight loss been absent, overall survival would likely have been longer. Consistent with this interpretation, the spleens of FHD-286–treated animals were significantly smaller than those of vehicle-treated controls, indicating a lower B-ALL burden (**Figure 4d**).

**Figure 4:**
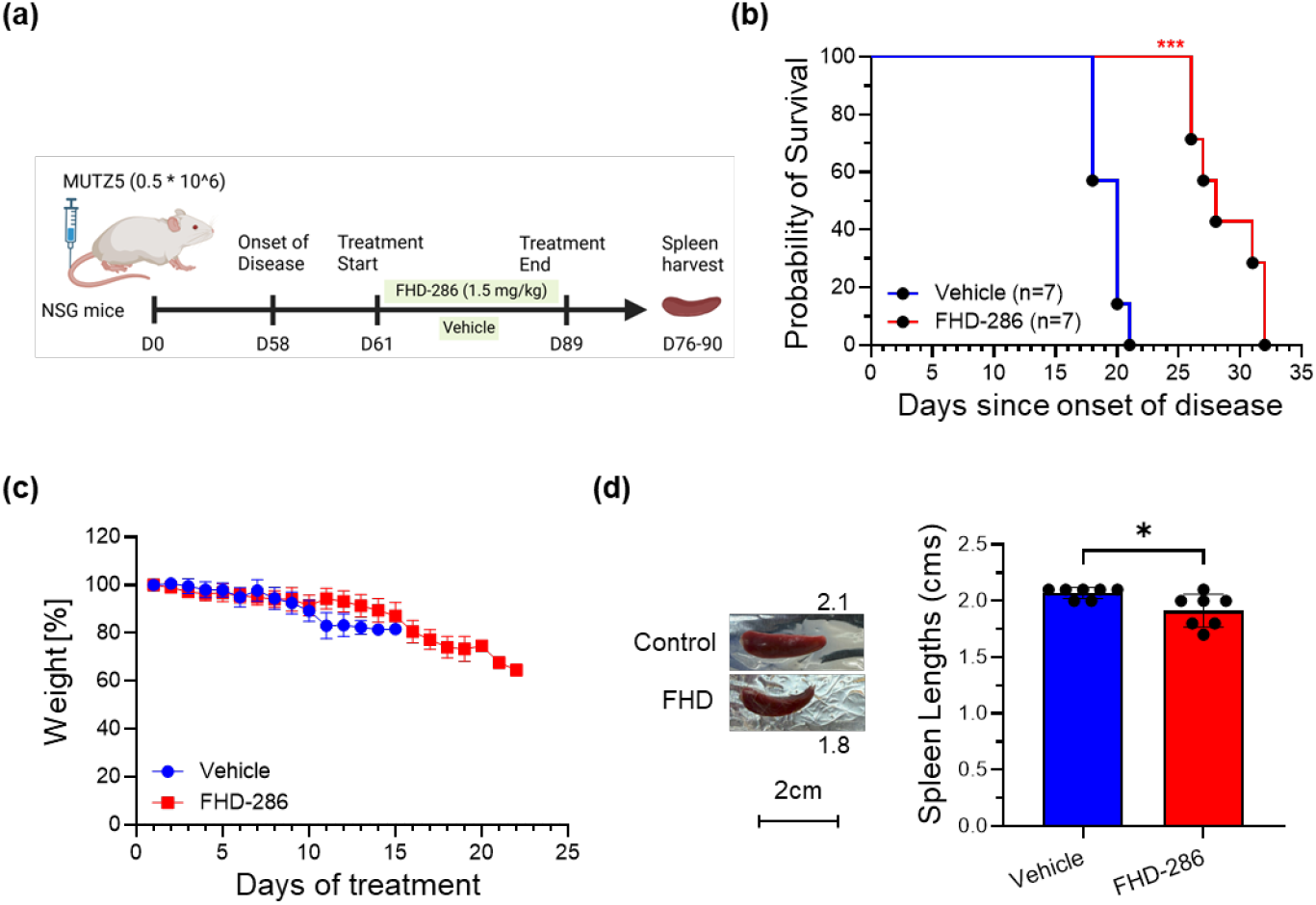
Pharmacologic Inhibition of BRG1 Extends the Survival of Ph-like B-ALL Model Mice. **A)** Schematic of the MUTZ5 xenograft model. NSG mice harboring MUTZ5 xenografts were treated with FHD-286 (1.5 mg/kg, via oral gavage) or vehicle continuously until the end of the study. **B**) Kaplan-Meier survival analysis of Ph-like PDX model (MUTZ5) mice treated for 22 days with vehicle control or FHD-286 (100 nM) (n= 7 mice/cohort). Significance was calculated via a log-rank test in Prism. Data are represented as individual values with mean ± SEM bars. *P < 0.05; **P < 0.01; ***P < 0.001 by t-test or Log-rank (Mantel-Cox) test. **C)** The weights of mice in different treatment groups were monitored throughout the treatment to test for drug-associated toxicities. **D)** The spleens of the mice were isolated after sacrifice, and spleen sizes were plotted for each treatment group. A representative image of a spleen from each treatment group is shown. Data are represented as individual values with mean ± SEM bars. *P < 0.05; **P < 0.01; ***P < 0.001 by two-tailed paired t-test.

In conclusion, these findings demonstrate that the pharmacologic inhibition of BRG1/BRM offers significant anti-leukemic activity in Ph-like ALL *in vitro* and *in vivo and* supports future investigation into the mechanistic underpinnings of BRG1-mediated leukemic survival.

## Discussion

BRG1, encoded by *SMARCA4*, is a core ATPase subunit of the SWI/SNF (BAF) chromatin-remodeling complex and plays a well-established role in regulating chromatin accessibility and gene transcription during hematopoietic development. In early B cell development, BRG1 is particularly crucial during the pro-to pre-B cell transition, facilitating lineage-specific gene expression programs. While *SMARCA4* has been extensively studied in solid tumors and in Burkitt lymphoma - where loss-of-function mutations are frequently associated with poor prognosis and support its tumor suppressor role^24-26,29^ - its function in B-cell acute lymphoblastic leukemia (B-ALL) remains poorly characterized.

Our genomic and functional analyses reveal a striking absence of inactivating *SMARCA4* mutations in B-ALL, suggesting that BRG1 function is preserved in this context. Likewise, no recurrent or functionally significant *SMARCA2* mutations were identified in large B-ALL cohorts. By contrast, activating NRAS and KRAS mutations are observed in over 30% of B-ALL cases, consistent with previous studies^30^. Using genome-wide CRISPR loss-of-function screening data (DepMAP), we find that *SMARCA4* is essential for the survival of B-ALL cells. Notably, among B-cell-derived malignancies, B-ALL exhibits the highest dependency on *SMARCA4*, whereas *SMARCA2* appears largely dispensable. However, this dependency varies across B-ALL cell lines, suggesting that BRG1 may represent a subtype-specific vulnerability.

By integrating gene expression data from the COG P9906 clinical trial, we identified that Ph-like B-ALL, an aggressive, kinase-driven subtype, exhibits significantly higher *SMARCA4* expression compared to *KMT2A*-R B-ALL. Importantly, high *SMARCA4* or *SMARCA2* expression correlated with inferior overall survival in Ph-like B-ALL, while higher *SMARCA4* expression predicted improved outcomes in *KMT2A*-R patients. These context-dependent associations suggest a potential oncogenic role for BRG1 in Ph-like B-ALL, in contrast to a tumor-suppressive role in other subtypes, such as those with KMT2A fusions.

Our findings support the emerging paradigm that chromatin regulators like BRG1 can function as oncogenes or tumor suppressors depending on the transcriptional and epigenetic context. In Ph-like B-ALL, BRG1 may cooperate with activated kinase signaling pathways (e.g., JAK-STAT, ABL-class fusions) to maintain oncogenic gene expression programs. Our data align with prior findings that Ph-like B-ALL is highly transcriptionally active and epigenetically plastic, making it potentially more reliant on chromatin remodeling factors^3^.

Functionally, BRG1 inhibition using BRM014 and FHD-286 induced potent G0/G1 arrest in Ph-like B-ALL cell lines, consistent with a role in regulating cell-cycle progression. These effects mirror observations in the RN2 AML model, where BRG1 depletion also led to cell-cycle arrest without inducing apoptosis^14^. In contrast, high-risk *KMT2A*-R B-ALL cells were less responsive to BRG1 inhibition, possibly due to compensatory chromatin remodeling driven by KMT2A fusion proteins^31^. Prior studies have shown that KMT2A fusions can recruit DOT1L and other epigenetic cofactors to drive leukemogenic transcription independently of BRG1^32-35^.

Although BRG1 inhibitors did not induce significant apoptosis *in vitro*, their cytostatic effects were sufficient to impair leukemia progression *in vivo*. Treatment of Ph-like B-ALL-engrafted *NSG* mice with FHD-286 significantly extended survival, providing strong preclinical support for BRG1 as a therapeutic target in this high-risk subtype. These findings are particularly compelling given the poor prognosis and limited targeted therapy options currently available for Ph-like B-ALL. Notably, even with advanced immunotherapeutic strategies such as CAR-T cell therapy followed by allogeneic hematopoietic stem cell transplantation, the estimated 3-year overall survival in patients with Ph-like ALL remains approximately 65%, underscoring the continued need for additional therapeutic approaches^36^.

The present work positions BRG1 inhibition as a rational therapeutic strategy for Ph-like B-ALL. Future studies should test whether an orally available BRG1 inhibitor can be “layered” onto standard multi-agent induction or consolidation regimens (vincristine, dexamethasone, anthracyclines, cytarabine, etc.) without creating scheduling conflicts or excess toxicity.

A second translational avenue is suggested by Battistello *et al*., who showed that a short *ex vivo* pulses of mSWI/SNF inhibitors during CAR-T manufacture reduced exhaustion markers and improved CAR-T function^37^. Building on those findings, it will be important to determine whether *in vivo* BRG1 inhibition, given concurrently with, or shortly after, CD19 CAR-T infusion, can further enhance CAR-T expansion, persistence, and cytotoxicity while simultaneously exerting a cytostatic brake on residual Ph-like leukemic cells.

In summary, we establish BRG1/SMARCA4 as a subtype-specific, targetable dependency in Ph-like B-ALL. The absence of inactivating mutations, the high oncogenic expression, the robust cytostatic response to pharmacologic inhibition, and the clear survival benefit *in vivo* collectively provide a compelling rationale to advance SMARCA4-directed therapies, either with front-line chemotherapy or in combination with CAR-T immunotherapy, into clinical testing for this high-risk leukemia.

## Data Availability

All data used in this study are publicly available and can be accessed from the Gene Expression Omnibus: GSE297161, GSE33315, GSE26408, GSE11877, and GSE38463.

## Acknowledgments

These studies were supported by the National Institutes of Health/National Cancer Institute (NIH/NCI) K22CA251649 (CH) and Alex’s Lemonade Stand Foundation (CH and AP). TS is supported by NIH/NCI R01CA186238. SS is supported by a Scholar Award from the St. Baldrick’s Foundation, a Leukemia Research Foundation New Investigator Award, and R37CA276517-01A1 from the National Cancer Institute of the National Institutes of Health. Graphical abstract is created in BioRender. Ayyadevara, A. (2025) https://BioRender.com/ckg8uh7

## Authorship Contributions

VSSAA planned and performed experiments, analyzed, and interpreted data, and wrote the manuscript; SG performed experiments and edited the manuscript; AP, RP, and MT performed experiments; TS assisted with study design; GW provided critical reagents; HG and SS performed computational analyses of gene expression datasets and survival studies; CH oversaw the study, designed and directed the study, analyzed, and interpreted data, provided research funding and experimental materials, edited the manuscript. All authors approved the final version of the manuscript.

## Materials and Methods

### Human ALL cell lines

Human *KMT2A*-R (SEM, HB11;19, RS4;11, KOPN8) and Ph-like (MUTZ5, MHH-CALL4) B-ALL cell lines listed in **Supplementary Table 2**, were obtained from the DSMZ biorepository (Braunschweig, Germany) validated by short tandem repeat (STR) analysis, and confirmed to be Mycoplasma-free every six months^38-41^. Cells were cultured in RPMI 1640 medium (Life Technologies) supplemented with GlutaMAX, 20% fetal bovine serum (FBS), 100 IU/mL penicillin, and 100 μg/mL streptomycin (hereafter referred to as “B cell medium”), at 37 °C in a humidified incubator with 5% CO_2_. Cell lines were maintained in culture for fewer than three months to minimize the risk of infectious contamination and genetic drift.

### Flow Cytometry Analysis

Human leukemia samples were stained with cell surface antibodies (**Supplementary Table 3**) following the manufacturers’ instructions. Annexin V-PE (Invitrogen), 7-AAD (Invitrogen), or DAPI (BD Biosciences) were used for apoptosis analysis. Flow cytometry was performed using MACSQuant flow cytometers (Miltenyi), and data were analyzed with FlowJo software (TreeStar).

### *In Vivo* Experimentation

Eight-to ten-week-old female NSG (NOD.Cg-*Prkdc*^*scid*^ *Il2rg*^*tm1Wjl*^/SzJ) mice were obtained from Jackson Laboratory and housed under conditions approved by the IACUC of Loma Linda University. A total of 14 mice (7 per group) were injected with 0.5 million MUTZ5 cells via tail vein. Leukemia progression was monitored by collecting blood from the submandibular vein, lysing red blood cells with ACK buffer (Gibco), staining with FITC-conjugated anti-human CD45 and PerCP-Cy5.5-conjugated anti-human CD19, and analyzing by flow cytometry.

Once leukemia onset was confirmed (defined as 1–5% ALL cells in blood), FHD-286 was administered by oral gavage at 1.5 mg/kg daily until study endpoint. FHD-286 was prepared in a vehicle composed of 1.66% DMSO (Thermo Scientific), 40% PEG400 (MedChem Express), 5% Tween 80 (MedChem Express), and 53.34% PBS. Mice were euthanized upon reaching any of the following endpoint criteria: >80% CD19+/CD45+ cells in blood, ≥20% body weight loss, or impaired mobility due to disease progression or potential drug toxicity, as described previously^31,42-44^. Spleens were harvested, and ALL cells were isolated and cryopreserved.

### Cell Proliferation and Viability Assays

A total of 25,000 human ALL cells were seeded in 100 μL B cell medium per well of a 96-well Costar plate (Corning). BRM014 or FHD-286 (**Supplementary Table 4**) was diluted to the indicated concentrations in medium and added to a final culture volume of 150 μL. After 72 hours, cell proliferation and viability were assessed using the XTT assay (Cell Signaling), following the manufacturer’s protocol, with absorbance measured at 450 nm. Fold change was calculated relative to untreated controls (set to 100%).

### Western Blotting

Cells were lysed in CelLytic buffer (Sigma) supplemented with 1× HALT protease and phosphatase inhibitor cocktails (Thermo Scientific). Protein lysates were separated by SDS-PAGE on Bolt 4–12% Bis-Tris gradient gels (Invitrogen) and transferred to PVDF membranes (Millipore). Membranes were probed with primary antibodies (**Supplementary Table 3**) and corresponding HRP-conjugated or IRDye 680RD/800CW-conjugated secondary antibodies (Cell Signaling, LI-COR). Detection was performed using Amersham ECL reagents (GE Life Sciences).

### Quantitative Real-Time PCR (qRT-PCR)

Total RNA was extracted using the RNeasy kit (Qiagen), and cDNA was synthesized using the RevertAid First Strand cDNA Synthesis Kit (Thermo Scientific). Quantitative PCR was carried out with Power SYBR® Green Master Mix (Applied Biosystems) on a Stratagene Mx3005p instrument (Agilent), using standard cycling conditions. Primer sequences are listed in **Supplementary Table 5**.

### Statistical Analysis

Data visualization and statistical analyses were performed using GraphPad Prism 10. For *in vitro* studies, comparisons between two groups were made using unpaired two-tailed Student’s t-test. For comparisons involving three or more groups, one-way ANOVA with Dunnett’s or Sidak’s post hoc test was used. Murine survival analysis was performed using the log-rank (Mantel–Cox) test. A *P* value <0.05 was considered statistically significant. All results are reported as mean ± SEM.

### Declaration of Generative AI and AI-assisted Technologies in the Writing Process

During the preparation of this work the author(s) used ChatGPT in order to check grammar and punctuation. After using this tool/service, the author(s) reviewed and edited the content as needed and take(s) full responsibility for the content of the publication.

## References

1 Roberts, K. G. et al. Genetic alterations activating kinase and cytokine receptor signaling in high-risk acute lymphoblastic leukemia. Cancer Cell 22, 153–166 (2012). 10.1016/j.ccr.2012.06.005

2 Loh, M. L. et al. Tyrosine kinome sequencing of pediatric acute lymphoblastic leukemia: a report from the Children’s Oncology Group TARGET Project. Blood 121, 485–488 (2013). 10.1182/blood-2012-04-422691

3 Roberts, K. G. et al. Targetable kinase-activating lesions in Ph-like acute lymphoblastic leukemia. N Engl J Med 371, 1005–1015 (2014). 10.1056/NEJMoa1403088

4 Tasian, S. K. et al. High incidence of Philadelphia chromosome-like acute lymphoblastic leukemia in older adults with B-ALL. Leukemia 31, 981–984 (2017). 10.1038/leu.2016.375

5 Roberts, K. G. et al. High Frequency and Poor Outcome of Philadelphia Chromosome-Like Acute Lymphoblastic Leukemia in Adults. J Clin Oncol 35, 394–401 (2017). 10.1200/JCO.2016.69.0073

6 Herold, T. et al. Adults with Philadelphia chromosome-like acute lymphoblastic leukemia frequently have IGH-CRLF2 and JAK2 mutations, persistence of minimal residual disease and poor prognosis. Haematologica 102, 130–138 (2017). 10.3324/haematol.2015.136366

7 Jain, N. et al. Ph-like acute lymphoblastic leukemia: a high-risk subtype in adults. Blood 129, 572–581 (2017). 10.1182/blood-2016-07-726588

8 Den Boer, M. L. et al. A subtype of childhood acute lymphoblastic leukaemia with poor treatment outcome: a genome-wide classification study. The Lancet. Oncology 10, 125–134 (2009). 10.1016/S1470-2045(08)70339-5

9 Mullighan, C. G. et al. Deletion of IKZF1 and prognosis in acute lymphoblastic leukemia. N Engl J Med 360, 470–480 (2009). 10.1056/NEJMoa0808253

10 Roberts, K. G. & Mullighan, C. G. Genomics in acute lymphoblastic leukaemia: insights and treatment implications. Nat Rev Clin Oncol 12, 344–357 (2015). 10.1038/nrclinonc.2015.38

11 Harvey, R. C. et al. Rearrangement of CRLF2 is associated with mutation of JAK kinases, alteration of IKZF1, Hispanic/Latino ethnicity, and a poor outcome in pediatric B-progenitor acute lymphoblastic leukemia. Blood (2010). https://doi.org/blood-2009-09-245944 [pii] 10.1182/blood-2009-09-245944

12 Wilson, B. G. & Roberts, C. W. SWI/SNF nucleosome remodellers and cancer. Nat Rev Cancer 11, 481–492 (2011). 10.1038/nrc3068

13 Mittal, P. & Roberts, C. W. M. The SWI/SNF complex in cancer - biology, biomarkers and therapy. Nat Rev Clin Oncol 17, 435–448 (2020). 10.1038/s41571-020-0357-3

14 Shi, J. et al. Role of SWI/SNF in acute leukemia maintenance and enhancer-mediated Myc regulation. Genes Dev 27, 2648–2662 (2013). 10.1101/gad.232710.113

15 Papillon, J. P. N. et al. Discovery of Orally Active Inhibitors of Brahma Homolog (BRM)/SMARCA2 ATPase Activity for the Treatment of Brahma Related Gene 1 (BRG1)/SMARCA4-Mutant Cancers. J Med Chem 61, 10155–10172 (2018). 10.1021/acs.jmedchem.8b01318

16 Farnaby, W. et al. BAF complex vulnerabilities in cancer demonstrated via structure-based PROTAC design. Nat Chem Biol 15, 672–680 (2019). 10.1038/s41589-019-0294-6

17 Bossen, C. et al. The chromatin remodeler Brg1 activates enhancer repertoires to establish B cell identity and modulate cell growth. Nature Immunology 16, 775–784 (2015). 10.1038/ni.3170

18 Fischer, U. et al. Cell Fate Decisions: The Role of Transcription Factors in Early B-cell Development and Leukemia. Blood Cancer Discov 1, 224–233 (2020). 10.1158/2643-3230.BCD-20-0011

19 Holmfeldt, L. et al. The genomic landscape of hypodiploid acute lymphoblastic leukemia. Nature Genetics 45, 242–252 (2013). 10.1038/ng.2532

20 Choi, J. et al. The SWI/SNF-like BAF Complex Is Essential for Early B Cell Development. The Journal of Immunology 188, 3791–3803 (2012). 10.4049/jimmunol.1103390

21 Green, M. R. et al. Signatures of murine B-cell development implicate Yy1 as a regulator of the germinal center-specific program. Proceedings of the National Academy of Sciences 108, 2873–2878 (2011). 10.1073/pnas.1019537108

22 Zhang, J. et al. The genetic basis of early T-cell precursor acute lymphoblastic leukaemia. Nature 481, 157–163 (2012). 10.1038/nature10725

23 Bowman, W. P. et al. Augmented therapy improves outcome for pediatric high risk acute lymphocytic leukemia: Results of Children’s Oncology Group trial P9906. Pediatric Blood & Cancer 57, 569–577 (2011). 10.1002/pbc.22944

24 Tian, Y., Xu, L., Li, X., Li, H. & Zhao, M. SMARCA4: Current status and future perspectives in non-small-cell lung cancer. Cancer Lett 554, 216022 (2023). 10.1016/j.canlet.2022.216022

25 Pang, L. L. et al. SWI/SNF family mutations in advanced NSCLC: genetic characteristics and immune checkpoint inhibitors’ therapeutic implication. ESMO Open 9, 103472 (2024). 10.1016/j.esmoop.2024.103472

26 Orvis, T. et al. BRG1/SMARCA4 inactivation promotes non-small cell lung cancer aggressiveness by altering chromatin organization. Cancer Res 74, 6486–6498 (2014). 10.1158/0008-5472.CAN-14-0061

27 Brady, S. W. et al. The genomic landscape of pediatric acute lymphoblastic leukemia. Nat Genet 54, 1376–1389 (2022). 10.1038/s41588-022-01159-z

28 Fiskus, W. et al. BRG1/BRM inhibitor targets AML stem cells and exerts superior preclinical efficacy combined with BET or menin inhibitor. Blood 143, 2059–2072 (2024). 10.1182/blood.2023022832

29 Deng, Q. et al. SMARCA4 is a haploinsufficient B cell lymphoma tumor suppressor that fine-tunes centrocyte cell fate decisions. Cancer Cell 42, 605-622.e611 (2024). 10.1016/j.ccell.2024.02.011

30 Chan, L. N. et al. Signalling input from divergent pathways subverts B cell transformation. Nature 583, 845–851 (2020). 10.1038/s41586-020-2513-4

31 Ayyadevara, V. et al. DYRK1A inhibition results in MYC and ERK activation rendering KMT2A-R acute lymphoblastic leukemia cells sensitive to BCL2 inhibition. Leukemia (2025). 10.1038/s41375-025-02575-w

32 Bernt, M., Kathrin et al. MLL-Rearranged Leukemia Is Dependent on Aberrant H3K79 Methylation by DOT1L. Cancer Cell 20, 66–78 (2011). 10.1016/j.ccr.2011.06.010

33 Dafflon, C. et al. Complementary activities of DOT1L and Menin inhibitors in MLL-rearranged leukemia. Leukemia 31, 1269–1277 (2017). 10.1038/leu.2016.327

34 Krivtsov, A. V. et al. A Menin-MLL Inhibitor Induces Specific Chromatin Changes and Eradicates Disease in Models of MLL-Rearranged Leukemia. Cancer Cell 36, 660–673 e611 (2019). 10.1016/j.ccell.2019.11.001

35 Dafflon, C., Tiedt, R. & Schwaller, J. Targeting multiple nodes of MLL complexes to improve leukemia therapy. Oncotarget 8, 90614–90615 (2017). 10.18632/oncotarget.21598

36 Dai, H. P. et al. CAR-T cell therapy followed by allogenic hematopoietic stem cell transplantation yielded comparable outcome between Ph like ALL and other high-risk ALL. Biomark Res 11, 19 (2023). 10.1186/s40364-023-00451-2

37 Battistello, E. et al. Stepwise activities of mSWI/SNF family chromatin remodeling complexes direct T cell activation and exhaustion. Mol Cell 83, 1216–1236 e1212 (2023). 10.1016/j.molcel.2023.02.026

38 Gotesman, M. et al. mTOR inhibition enhances efficacy of dasatinib in ABL-rearranged Ph-like B-ALL. Oncotarget 9, 6562–6571 (2018). 10.18632/oncotarget.24020

39 Surrey, L. F. et al. Clinical utility of custom-designed NGS panel testing in pediatric tumors. Genome Med 11, 32 (2019). 10.1186/s13073-019-0644-8

40 Joseph, P. L. et al. Combinatorial efficacy of entospletinib and chemotherapy in patient-derived xenograft models of infant acute lymphoblastic leukemia. Haematologica 106, 1067–1078 (2020). 10.3324/haematol.2019.241729

41 Hurtz, C. et al. Oncogene-independent BCR-like signaling adaptation confers drug resistance in Ph-like ALL. The Journal of Clinical Investigation 130, 3637–3653 (2020). 10.1172/JCI134424

42 Hurtz, C. et al. Redundant JAK, SRC and PI3 Kinase Signaling Pathways Regulate Cell Survival in Human Ph-like ALL Cell Lines and Primary Cells. Blood 130, 717–717 (2017).

43 Kruth, K. A. et al. Suppression of B-cell development genes is key to glucocorticoid efficacy in treatment of acute lymphoblastic leukemia. Blood 129, 3000–3008 (2017). 10.1182/blood-2017-02-766204

44 Tasian, S. K. et al. Potent efficacy of combined PI3K/mTOR and JAK or ABL inhibition in murine xenograft models of Ph-like acute lymphoblastic leukemia. Blood 129, 177–187 (2017). 10.1182/blood-2016-05-707653

