## Supplementary Information for "Selective Sensitivity of Ph-like B-ALL to BRG1 Inhibition Reveals a Novel Targeted Therapy Strategy"

**Supplementary Figures:**

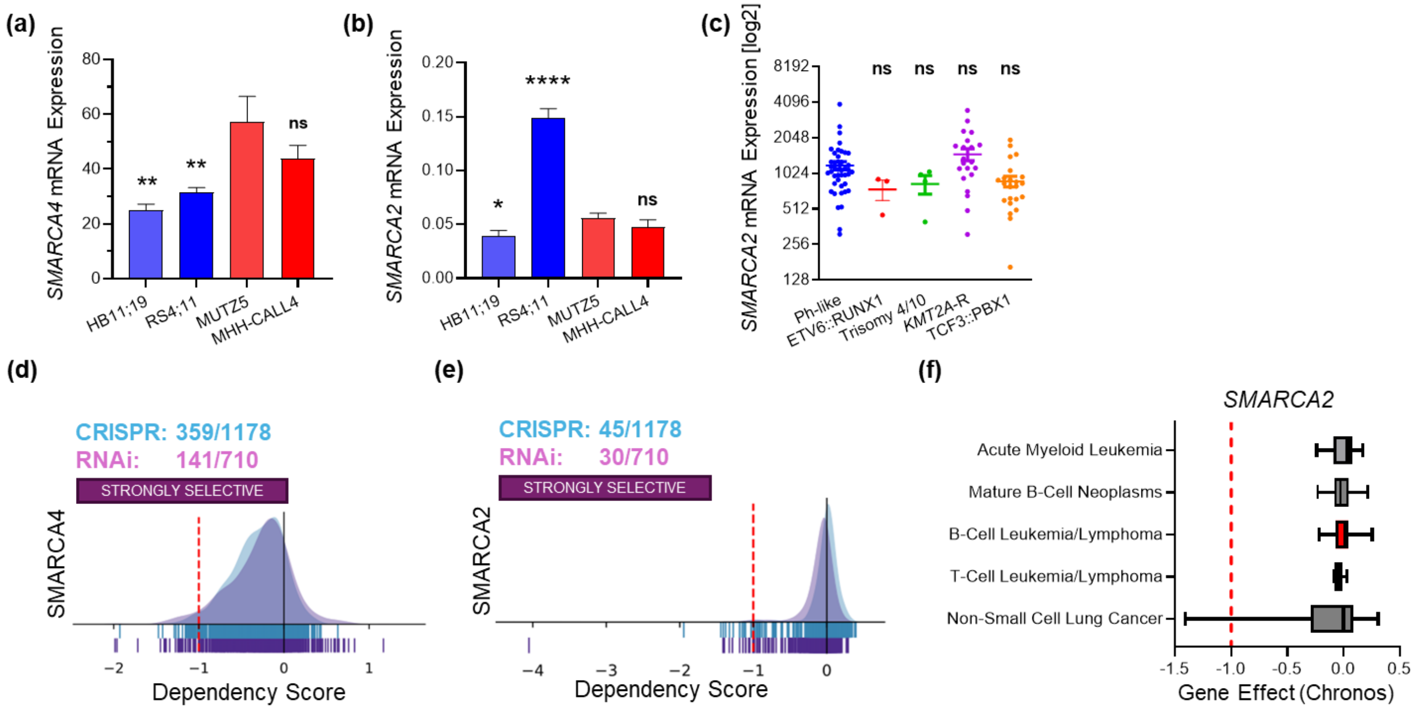


**Supplementary Figure 1:** RT-PCR was performed to determine the mRNA expression levels of the *SMARCA4* **(A)** and *SMARCA2* **(B)** in Ph-like (MUTZ5 and MHH-CALL4) and *KMT2A*-R B-ALL cell lines (HB11;19 and KOPN8) after 24 hours of treatment either with control or 100 nM FHD-286. *ACTB* (Beta Actin) was used as a reference gene to calculate expression levels. **C)** mRNA Expression of *SMARCA2* in different subtypes of B-ALL populations (GSE11877). Dependency analysis of *SMARCA4* **(D)** and *SMARCA2* **(E, F)** via the Cancer Dependency MAP website. Data are represented as individual values with mean ± SEM bars. *P < 0.05; **P < 0.01; ***P < 0.001; ****P < 0.0001 by One-way Anova Dunnett's multiple comparison test.

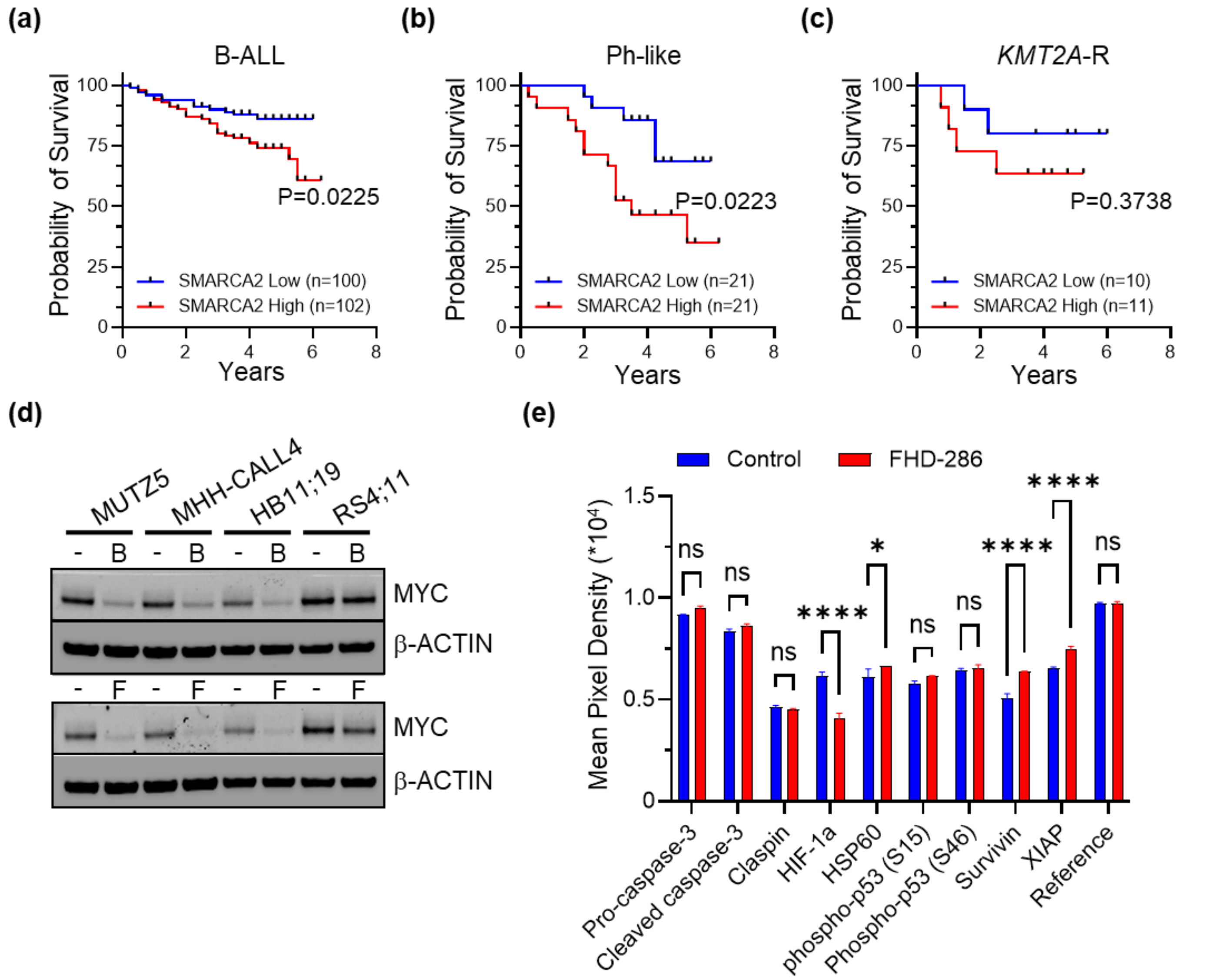


**Supplementary Figure 2:** Kaplan-Meier analysis of the overall survival of all B-ALL patients **(A)**, Ph-like B-ALL **(B)**, and *KMT2A*-R B-ALL **(C)** patients from the COG study P9906 with high vs low *SMARCA2* mRNA expression (GSE11877). D**)** Western blot analysis of MYC and β-ACTIN in two *KMT2A*-R (HB11;19, RS4;11) and two Ph-like (MUTZ5, MHH-CALL4) B-ALL cell lines treated with control or BRM014 (100 nM) or FHD-286 (100 nM) for 24 hours. **G)** Analysis of changes in the expression of an array of apoptosis-related genes in Ph-like (MUTZ5) B-ALL cell line treated with control or 100 nM FHD-286 for 24 hours. Significance was calculated via log-rank test in Prism **(A-C)**. Data are represented as individual values with mean ± SEM bars. *P < 0.05; **P < 0.01; ***P < 0.001; ****P < 0.0001 by Two-way Anova Sidak's multiple comparison test **(E)** or Two-sided log-rank (Mantel-Cox) test **(A-C)**.

**Supplementary Table 1: Details of *SMARCA4* mutations in B-cell patients.**

| # | Patient identifier | ALL subtype | Chromosome | Position (hg19) | Reference allele | Mutated allele | Mutation | Amino acid change | Mutation type |
|---|---|---|---|---|---|---|---|---|---|
| **1** | SJBALL004114 | Ph-like_CRLF2 | chr19 | 11129638 | T | C | Non-synonymous | L815P | missense |
| **2** | SJBALL022167 | B-other | chr19 | 11144113 | G | A | Non-synonymous | G1232S | missense |
| **3** | SJCOGALL010219 | Ph-like_CRLF2 | chr19 | 11144798 | G | A | Non-synonymous | E1292_E28splice | splice |
| **4** | SJCOGALL011148 | Hyperdiploid | chr19 | 11144135 | CC | TT | Non-synonymous | S1239F | missense |
| **5** | SJCOGALL011164 | B-other | chr19 | 11134231 | G | C | Non-synonymous | R966P | missense |
| **6** | SJHYPER141 | Hyperdiploid | chr19 | 11144146 | C | T | Non-synonymous | R1243W | missense |
| 7 | PANZXC | B-other | chr19 | 11097202 | C | A | Synonymous | G231G | silent |
| | | | chr19 | 11097205 | T | C | Synonymous | P232P | silent |
| 8 | SJALL016639 | ETV6-RUNX1 | chr19 | 11113696 | C | G | Synonymous | P605_E12 splice region | splice region |

**Supplementary Table 2: Leukemia cell lines used in these studies.**

| Cell Line | Lesion | Leukemia type | Source |
|---|---|---|---|
| **SEM** | *KMT2A::AFF1* | B-ALL | DSMZ |
| **RS4;11** | *KMT2A::AFF1* | B-ALL | DSMZ |
| **HB11;19** | *KMT2A::MLLT1* | B-ALL | Dr. Sarah Tasian, CHOP |
| **KOPN8** | *KMT2A::MLLT1* | B-ALL | DSMZ |
| **MUTZ5** | *IGH::CRLF2* | B-ALL | DSMZ |
| **MHH-CALL4** | *IGH::CRLF2* | B-ALL | DSMZ |

**Supplementary Table 3: Antibodies used in these studies.**

***Western blotting***

| Antigen | Clone | Manufacturer | ID # |
|---|---|---|---|
| **BRG1** | D1Q7F | Cell signaling | 49360S |
| **BRM** | D9E8B | Cell signaling | 11966S |
| **MYC** | D84C12 | Cell signaling | 5605S |
| **Beta-Actin** | 13E5 | Cell signaling | 4970L |

***Flow cytometry***

| Antigen | Clone | Manufacturer | ID # |
|---|---|---|---|
| **CD19** | HIB19 | Invitrogen | 45-0199-42 |
| **CD45** | HI30 | Invitrogen | 11-0459-42 |

**Supplementary Table 4: Targeted inhibitors used in these studies.**

| Name | Vendor | ID # |
|---|---|---|
| **BRM/BRG1 ATP Inhibitor-1 (Compound 14) or BRM014** | Selleckchem | E0111 |
| **FHD-286** | MedChem express | HY-144835 |

**Supplementary Table 5: RT-PCR primers used in these studies.**

| # | Gene | Forward primer (5' to 3') | Reverse primer (5' to 3') |
|---|---|---|---|
| 1 | **SMARCA4** | TGG CCT GCA GTC CTA CTA T | GCG AGG ATG TGC TTG TCT TT |
| 2 | **SMARCA2** | GGG CTT GGA AAG ACC ATA CA | CCA GAG TTC AGG GAG CTT ATT C |
| 3 | **CCND3** | CTT ACT GGA TGC TGG AGG TAT G | CGG GTA CAT GGC AAA GGT ATA A |
| 4 | **CDK4** | CAG TTC GTG AGG TGG CTT TA | GCC ATC TGG TAG CTG TAG ATT C |
| 5 | **CDK6** | ACC AGC AGC GGA CAA ATA A | CAA CAT CTC TAG GCC AGT CTT C |
| 6 | **E2F1** | TGC AGA GCA GAT GGT TAT GG | GAT GGT GGT GGT GAC ACT ATG |
| 7 | **MYC** | AGG GAG ATC CGG AGC GAA TA | GTC CTT GCT CGG GTG TTG TA |
| 9 | **CDKN1B** | GTC AAA CGT GCG AGT GTC TA | TGC AGG TCG CTT CCT TAT TC |
| 10 | **ACTB** | ACA GAG CCT CGC CTT TG | CCT TGC ACA TGC CGG AG |
